# Accurate Cell Abundance Quantification using Multi-positive and Unlabeled Self-learning

**DOI:** 10.1101/2024.10.12.617956

**Authors:** Yating Lin, Xiaoqi Chen, Yuxiang Lin, Xu Xiao, Zhibin Huang, Wenxian Yang, Rongshan Yu, Jiahuai Han

## Abstract

Quantifying the abundance of different cell types in pathological samples can help to uncover the correlations between cell composition and pathological conditions, offering deeper insights into the roles of different cell types in complex diseases. Conventional methods for cell abundance estimation often employ unsupervised clustering or supervised learning to identify cell types and estimate their proportions. However, these methods face challenges in accurately quantifying cell abundances, as clustering results could be unreliable and supervised methods may misclassify cell types not presented in the training data. We introduce Clever^XMBD1^ (denoted as Clever) for quantifying cell abundance from complex samples using a multi-class classifier trained with a confidence-based multi-positive and unlabeled loss function. Our evaluations show that Clever consistently and substantially outperforms existing methods in quantifying cell type abundance across multiple single cell datasets derived from different modalities, including CyTOF and image mass cytometry.

## Introduction

High-throughput cellular profiling methods, such as single-cell multiplexed imaging ^1^ and mass cytometry ^2^, enable the measurement of multiple marker expressions at unprecedented resolutions in individual cells. These techniques pave the way for in-depth investigations of cellular interations within living organisms and their associations with pathological conditions ^3–5^. Recently, an increasing number of single-cell studies have shifted their focus from qualitative analysis towards the quantitative analysis of cell abundance under various experimental or pathological conditions, with the aim of identifying the roles of different cell types in complex diseases ^6, 7^. In such context, the ability to derive accurate abundances of different cell populations becomes vital.

*Cluster-then-annotate approaches* have been extensively adopted in single-cell experiments ^8–11^. These methods start by clustering cells into distinct groups using unsupervised clustering. Then the cell types of the clusters can be identified by matching their expression patterns of a predefined set of markers or signatures to those of established cell types, and the abundance of each individual cell type can be derived. However, cell type abundances obtained by such methods can be unstable as the results of clustering algorithms are often highly sensitive to the adjustable parameters. In addition, the interpretation of clusters can be heuristic and lacks reproducibility, resulting in inconsistent cell type abundance quantifications across different experiments and research groups. Lastly, when analyzing large-scale datasets, some clustering algorithms may suffer from information loss due to random sub-sampling ^12^, which can be particularly problematic for identifying non-canonical or rare cell types ^13^. These intrinsic limitations of unsupervised clustering may lead to unreliable and inconsistent estimations of cell type abundance, potentially confounding the final biological analysis outcomes.

Alternatively, *marker-based* or *semi-supervised methods* assign cells to distinct cell types based on markers of known cell types either through an explicit gating strategy or an implicit statistical learning process. Gating assigns individual cells to discrete cell types based on hierarchically arranged bi-axial plots ^14^. In practice, cell type labels derived from gating procedures have been designated as silver standards due to the lack of groundtruth ^15^. However, cells distributed near or on the gating boundaries may not be assigned to any cell type after manual gating, resulting in some unlabeled cells. Excluding these unlabeled cells leads to an underestimation of cell type abundance, potentially causing erroneous interpretation of crucial cell types. Besides manual gating, statistical learning methods ^16, 17^ have also been developed to integrate marker information and infer cell types. However, these methods are highly sensitive to marker selection, where incorrect or incomplete markers can lead to inaccurate cell type identification and quantification.

*Supervised methods* use machine learning models trained on a reference dataset to derive cell type abundances ^18–20^. They do not specify markers explicitly but leverage on machine learning models to identify complex patterns and relationships within the data. By utilizing large annotated datasets, these approaches can potentially achieve higher accuracy and reproducibility in cell type annotation and abundance estimation. However, they encounter difficulties when dealing with unannotated or novel cell types not present in the training data, which could result in misclassifications and reduced generalizability. The lack of a robust mechanism to address these unknown cell types can ultimately distort downstream analysis results ^19^. Additionally, supervised methods are prone to overfitting to the preferences of the human annotator ^17^, as their performance heavily depends on manual gating, which can be unstable due to the manual selection of gating thresholds.

To address these challenges, we developed Clever (CelL abundancE quantification using multi-positiVe unlabEled leaRning), a semi-automated, deep learning-based computational framework to effectively identify and quantify cells from single-cell multiplexed imaging and proteomic datasets. Clever addresses the limitations of manual cell-type gating by considering the gated results as a multi-positive and unlabeled (MPU) dataset, where cells confidently identified through gating are treated as labeled cells with respective positive cell types, and the remaining cells are considered as unlabeled cells. Using an MPU strategy, Clever assigns the unlabeled cells into either one of the labeled cell types or an “Unknown cells” class if they belong to novel cell types not captured by gating. Moreover, Clever utilizes self-training to iteratively refine the classification of unlabeled cells, thereby reducing the dependence of the classification results on the positive cell selection and mitigating the effects of human biases in positive cell labelling. Benchmarking on various public image mass cytometry (IMC) and CyTOF datasets demonstrated that Clever outperforms state-of-the-art methods in both cell type identification and abundance quantification, and provides robust results in idenfifying cell types with significant abundance differentce in single-cell experiments.

## Methods and materials

### The Clever workflow

The workflow of Clever consists of three main steps (Figure 1). First, manual gating with a predefined gating strategy is applied to separate the input data into subsets of labeled cells, and a subset of ungated (or unlabeled) cells. Next, a deep neural network (DNN)-based model is used to perform multi-class cell identification using a multi-positive and unlabeled (MPU) learning strategy on both labeled and unlabeled cells. Clever adopts an iterative self-training approach. During each iteration, the model adds unlabeled cells with high-confidence predictions to the labeled set when the model reaches a state of relative convergence. Cells with low-confidence predictions remain unlabeled. The model then iteratively trains on the updated dataset until no additional high-confidence predictions can be identified. In most real-world scenarios, the unlabeled cells often contain a significant proportion of cells from positive cell types, which can cause the trained model to overfit to the training data. To mitigate this issue, we used a bagging approach in training ^21^. Specifically, we randomly split the unlabeled cells into *k* subsets, and combine each subset with the labeled cells to form *k* training datasets. We then performed the above self-training process on these *k* training datasets to train *k* prediction models so that each model was trained only on a subset of unlabeled cells of positive cell types. Subsequently, the cell types of all unlabeled cells were predicted based on the aggregated results from the *k* trained models.

**Figure 1:**
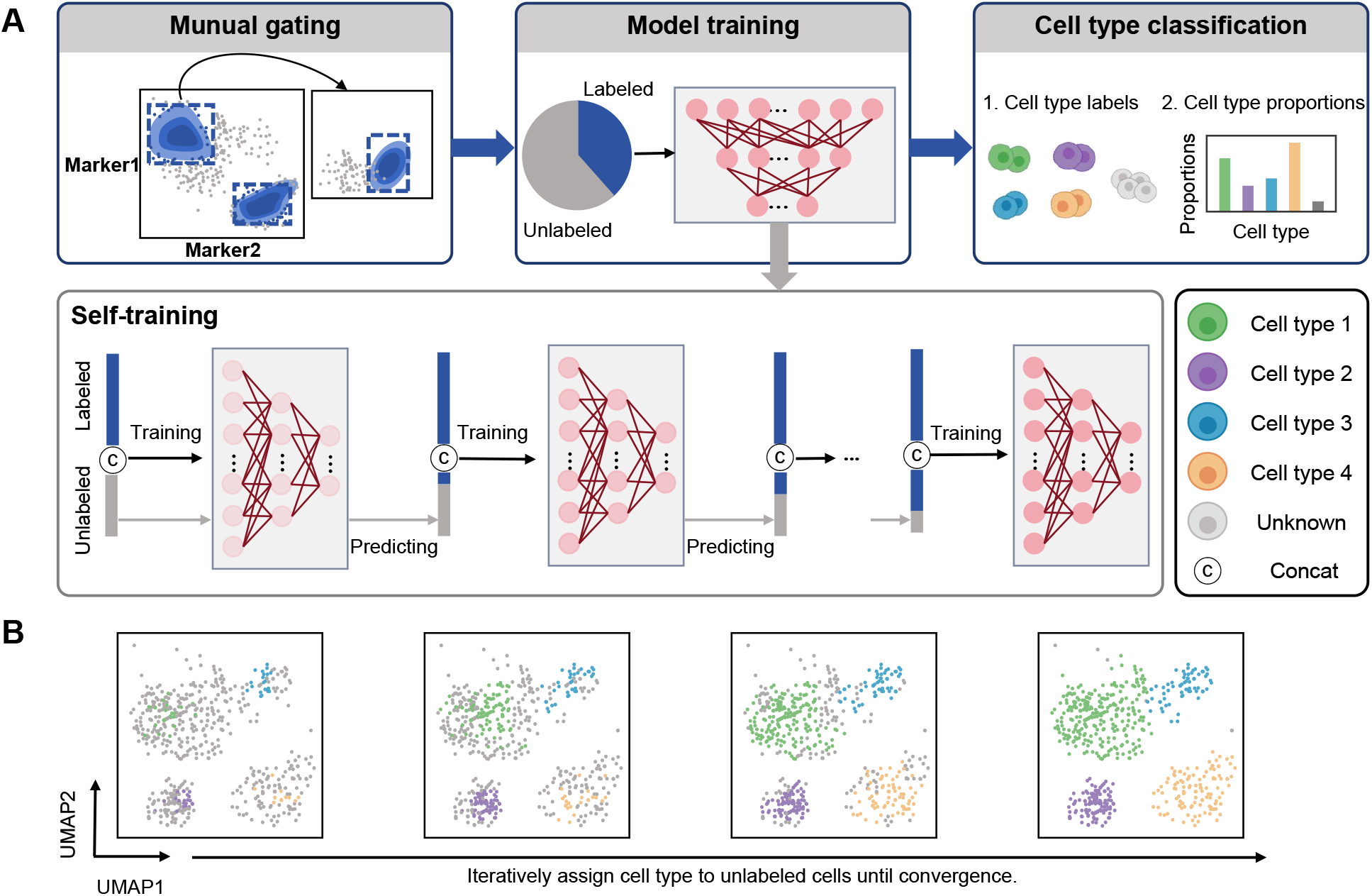
Clever workflow. (A) The Clever workflow consists of three main steps: (1) The initial step creates a training set by manual gating, resulting in a set of labeled cells with pre-identified cell types (labeled data *P*) and a set of unlabeled cells (unlabled data *U*). (2) The neural network model is trained using a self-training framework that iteratively leverages both labeled data and unlabeled data to minimize the Conf-MPU loss. (3) The trained models are applied to previous unlabeled data for cell type prediction. (B) Schematic diagram illustrating the iterative process in Clever’s cell type assignment to unlabeled cells.

#### Data preparation

We first use manual gating to separate the single cell dataset into *m* sets with known cell types, and one set of unlabled cells. Let *n*_P_ and *n*_U_ denote the number of labeled and unlabeled cells after gating, respectively. The entire dataset can be represented as a collection of labeled cells *P* and a collection of unlabeled cells *U* as follows:

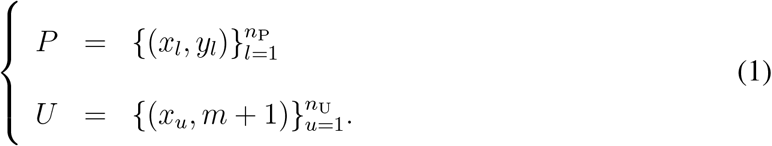

Here, 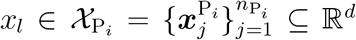 denotes the expressions of each labeled positive cell, where 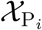 is drawn from 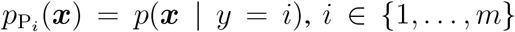, and *d* is the number of markers. *y*_*l*_ ∈ 𝒴_P_ = {1, 2, …, *m*} is the respective cell type for each labeled cell. For the unlabeled cell, 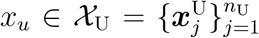 is assigned to the class label *y* = *m* + 1, where 𝒳_U_ is drawn from *p*_U_(***x***). The complete dataset is then represented as 𝒳 = 𝒳_P_ ∪ *𝒳*_U_ with labels 𝒴 = 𝒴_P_ ∪ {*m* + 1}.

#### Building multi-positive and unlabeled learning models

We build an MPU model to learn a decision function *f* : 𝒳 → 𝒴 to assign each unlabeled cell to one of the *m* + 1 classes to minimize a confidence-based multi-class positive and unlabeled loss ℒ_Conf-MPU_(*f*), which is defined as

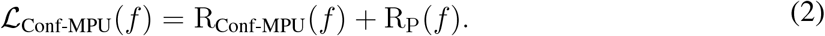

Here, R_Conf-MPU_(*f*) is the Conf-MPU risk estimator and R_P_(*f*) is a regularization term. R_Conf-MPU_(*f*) is defined as

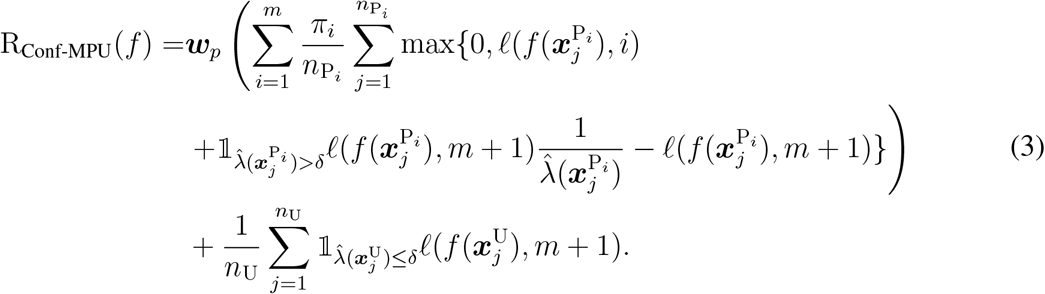

A detailed explanation of the risk function is provided in the Supplementary Information. In short, the loss function 𝓁() is implemented using the cross-entropy function. 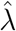serves as an empirical confidence score estimator. It is obtained by a classifier with a sigmoid output layer, ensuring that 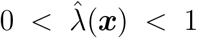. *δ* is the confidence threshold such that 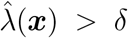 indicates that ***x*** is a positive sample. Otherwise, the sample is considered unknown. We set *δ* to 0.5 in this study. The empirical probability *π*_*i*_ is calculated based on the proportion of each positive class within individual training batches. The term ***w***_*p*_ represents the weight of positive risks, controlling the importance of positive classes relative to unknown classes. We set ***w***_*p*_ to 5 the first five training epochs to prioritize learning from positive samples, and reduce it to 1 afterwards to achieve a more balanced learning process and avoid overemphasizing the positive class.

The regularization term R_P_(*f*) leverages the KL-divergence to measure the difference between the model’s predictions and the assumed prior class distribution ^22^. This helps to prevent pre-mature convergence to suboptimal solutions and ensures more balanced learning across all classes, especially during the early stages of training. It is defined as

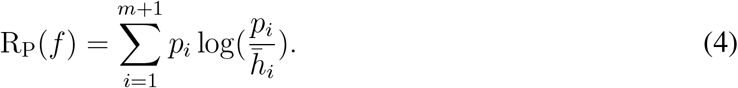

Specifically, we assume a uniform prior probability distribution *p*_*i*_ for each class *i*. 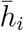 denotes the mean softmax probability for class *i* across all training samples, which is approximated within mini-batches.

#### Self-training process

A self-training strategy is employed to learn the decision function *f* : 𝒳 → 𝒴 and to expand the set of labeled cells by assigning pseudo-labels to cells in the unlabeled set 𝒳_U_. The MPU model is initially trained on the dataset 𝒳 = 𝒳_P_ ∪ 𝒳_U_ with labels 𝒴 = 𝒴_P_ ∪ {*m* + 1} until it reaches convergence. Here we define convergence as when the difference in training loss becomes less than 0.01 between consecutive epochs. Once this convergence is achieved, labels can be assigned to the unlabeled cells as follows.

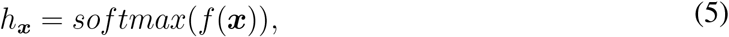

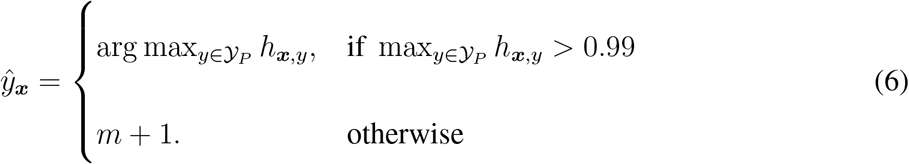

Here, *h*_***x***_ represents the probability distribution over the possible classes for an unlabeled cell ***x***, computed by applying the softmax function to the output of *f* (***x***). *h*_***x***,*y*_ denotes the predicted probability of cell ***x*** ∈ 𝒳_U_ being assigned to cell type *y*, where *y* ∈ 𝒴_P_. The pseudo-label 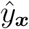 is assigned as the most confident label, corresponding to the cell type with the highest probability above 0.99. Otherwise, a default label *m*+1 is assigned to this cell. These pseudo-labels are treated as initial labels for the unlabeled cells, and cells with pseudo-labels are then incorporated into the labeled dataset for subsequent training. The process of predicting pseudo-labels and updating the labeled dataset is repeated iteratively until no additional high-confidence pseudo-labels can be generated for any unlabeled cells over five consecutive epochs.

### Implementation details and hyperparameters

We implemented our DNN model with PyTorch (v1.7.1) in Python (v3.8.18). The network consists of an input layer, fully connected hidden layers, and an output layer. It is trained using back-propagation with randomly initialized parameters. Training data were processed in mini-batches of 256 samples for each epoch. The Adam optimizer was employed with an initial learning rate of 0.05, which was reduced by a factor of 0.1 for every 30 epochs to promote convergence. To mitigate the risk of overfitting, a weight decay regularization term with a coefficient of 10^−5^ was applied. Training was conducted for a minimum of 30 epochs and stoped when there were no improvements in confident pseudo-label predictions over 5 consecutive epochs. The training procedure was conducted *k* times. The set of unlabeled cells *U* was randomly split into *k* equal-sized and non-overlapping subsets. Each subset was combined with the labeled cell set *P* to form the training data for one training process. After training, the final prediction for each cell in the original set *U* was obtained by averaging the probability outputs from all *k* models, where *k* was set to 5.

### Materials for benchmarking

#### Benchmark methods

We compared Clever with seven existing cell type identification methods including two marker-based methods, ACDC ^16^ and Astir ^17^, and five supervised methods, namely CyAnno ^19^, DGCyTOF ^20^, Random Forest, Support Vector Machine (SVM) and XGBoost. Among the benchmarking methods, ACDC, Astir, and CyAnno are capable of predicting unknown cell types. In all experiments, each method was provided with identical training and testing datasets, and executed using their respective recommended default parameters. Comprehensive descriptions and operational specifics of these methods are available in the Supplementary Information and Table S1.

#### Benchmark datasets

We used two IMC datasets and two CyTOF datasets for benchmarking. For the IMC datasets, the Hoch dataset ^23^ comprises data from 69 patients with metastatic melanoma, while the Mitamura dataset ^24^ includes skin punch biopsies from 12 COVID-19 patients and 4 healthy controls. For the CyTOF datasets, Levinedim13 ^25^ was generated from one healthy volunteer, comprising data of approximately 167,000 cells with 13 markers. Levinedim32 ^26^ was generated from two healthy human donors, consisting of around 266,000 cells with 32 protein-expression markers. Gating was used in these studies to generate cell type labels. The marker expression profiles of all live cells are transformed using arcsinh transformation with a co-factor set to the default value of 5. More details on the datasets are available in the Supplementary Information and Table S2.

### Cell annotation of unsupervised clustering methods

We employed Phenograph and FlowSOM for clustering. For cell annotation, we first standardize the log-count expression of all markers using the z-scaling method and averaged them for each identified cluster. We then computed the average expression of all markers for each specific cell type based on a predefined marker list. Clusters were assigned to the cell type with the highest average z-scaled value, while those associated with more than one cell type having similarly high values were categorized as “Unknown”.

### Simulated data generation

We evaluated the performance of the benchmark methods on synthetic datasets, which were generated using the approach developed by Kaspar et al ^27^. In particular, we define binary vectors for four cell types as

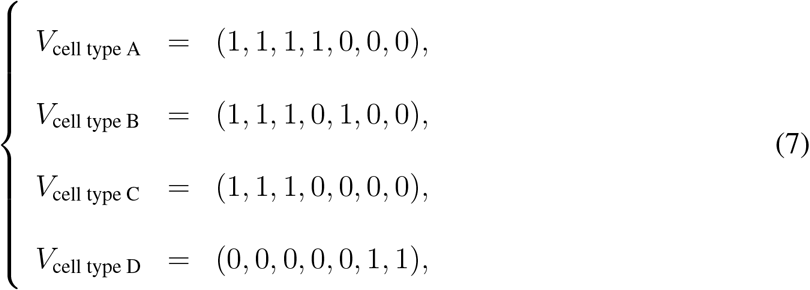

where ‘1’ and ‘0’ indicate high and low expressions of a specific marker, respectively. Using these cell type-specific marker vectors, we generate simulated data for each cell type with the following model:

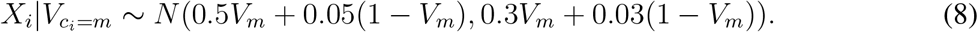

Here, *m* ∈ {*A, B, C, D*} represents different cell types. *X*_*i*_ follows a Gaussian distribution with the specified mean and variance values.

Our simulated dataset includes a case group and a control group, each consisting of 10 samples. To simulate variations in cell type abundance between the two groups, we deliberately selected the number of cells for each type to ensure no significant differences in the abundance of cell types A and B across the groups, while notable differences were observed in the abundance of cell types C and D. The average cell counts were set to (A: 2600, B: 2500, C: 2600, D: 400) for the case group, and (A: 2200, B: 2100, C: 1200, D: 1400) for the control group. To mimic potential biological variability, the number of each cell type in each sample varied within ± 350 from these baseline values.

### Simulated gating strategy

To investigate how the variations in manually selected gating thresholds for identifying labeled positive cells affect the annotation results in supervised methods, we generated additional training sets from the original labeled cells. We simulated the variations in gating thresholds by selecting cells with the highest 90%, 80%, 70%, 60%, and 50% expression levels of positive marker proteins among the initially labeled positive cell types. For cells characterized by negative markers, we chose those with the lowest 10%, 20%, 30%, 40%, and 50% expression levels of the corresponding negative marker proteins. Cells that did not meet the criteria for any specific cell type were designated as “unlabeled”. We conducted simulations using the Hoch dataset and defined cell types based on the following markers: B cells (CD20^+^ CD3^−^), BnT cells (CD20^+^ CD3^+^), CD4+ T cells (CD3^+^ CD4^+^ CD8^−^), CD8+ T cells (CD3^+^ CD8^+^ CD4^−^), FOXP3+ T cells (CD3^+^ FOXP3^+^), Macrophages (HLA-DR^+^ CD68^+^), Neutrophils (CD15^+^ MPO^+^), pDC (CD303^+^), Stroma (SMA^+^ Vimentin^+^), and Tumor (positive for any or multiple of SOX9, SOX10, MITF and S100A1). In the simulated dataset, we defined three cell types based on specific marker combinations: cell type, 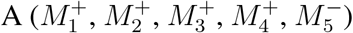 cell type 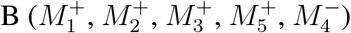, and cell type 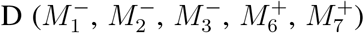. Here, *M*_*i*_ ∈ {*M*_1_, *M*_2_, *M*_3_, *M*_4_, *M*_5_, *M*_6_, *M*_7_} represents the markers as defined in the ‘Simulated Data Generation’ section, following their order in the binary vectors.

### Performance evaluation

We calculated a variety of metrics, including the F1 score, Cohen’s kappa, Matthews correlation coefficient, Adjusted Rand Index, sensitivity, and specificity, for the labeled results in each dataset using the labels from manual gating as the ground truth. For absolute cell abundance quantification, Lin’s Concordance Correlation Coefficient (CCC) and the Root-Mean-Square Error (RMSE) were used to assess the agreement between the predicted cell type proportions and the manual gating ground truth. To further enhance the robustness of our evaluation, we incorporated positive and negative controls. For positive controls, we assessed performance by measuring accuracy, i.e., the proportion of correctly assigned cells among the known cell types. For negative controls, we evaluated performance through the unassigned rate, representing the ratio of cells assigned as unknown class to all predicted unknown cells ^28^.

Additionally, to understand how various methods perform on imbalanced datasets, we quantify the imbalance level of each dataset as the Kullback-Leibler divergence between its training and testing set.

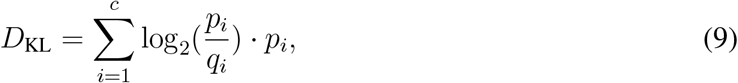

where *c* is the number of cell types, *p*_*i*_ and *q*_*i*_ represent the proportions of cell type *i* in the training set and the testing set, respectively.

## Statistics and reproducibility

All analysis are conducted using R (version 4.1.3) and Python (version 3.8). Statistical significance are determined using the Wilcoxon rank-sum test. All statistical tests used are defined in the figure legends.

## Results

### Unsupervised clustering leads to unstable cell abundance quantification

We first investigated how the adjustable parameter settings in unsupervised clustering methods would affect downstream cell identification and quantification results. We tested with two widely-used unsupervised clustering methods, Phenograph and FlowSOM, on Hoch dataset containing 10 pre-defined cell types ^24^. We applied each algorithm with various parameter settings, producing different numbers of output clusters. Specifically, for Phenograph, we varied the nearest neighbour parameter *k* in five increments, ranging from 20 to 100 (i.e., 20, 35, 50, 80, 100). For FlowSOM, we adjusted the cluster size parameter in five increments from 11 to 50 (Figures S1 and S2). Note that cluster sizes were chosen to match the expected number of cell types (e.g. 10 for the Hoch dataset) and varied such that potential cell states or “Unknown” cells could also be captured. Thus, we delibrately started from 11, resulting in the cluster sizes *k*= 11, 15, 20, 30 and 50.

Results indicated that changes in the number of clusters could lead to significant changes in the underlying cluster structure, as shown in Figure S1a. This result was highlighted by a uniform manifold approximation and projection, which demonstrated notable color intermixing indicative of changes in cluster delineations (Figure S1b). To assess the impact of different clustering results on cell type abundance quantification, we manually assigned cell types to each cluster (Methods). This process revealed that the merging and bifurcation of clusters led to significant fluctuations in the quantitative results following cell annotation (Figure S1c, d). In some parameter settings, certain cell types, such as CD8+ T and CD4+ T cells, were not successfully annotated. We further checked whether the abundances of distinct cell types would vary with different clustering parameters across various conditions. Our findings indicated that the results of cell type abundance were indeed dependent on the chosen clustering parameters. Although significant features in cell abundance were observed with certain parameter settings, these features could disappear when analyzed with alternative parameters.

We observed that with varying cluster structures obtained from different parameter settings in unsupervised clustering, downstream analysis may result in significantly different biological interpretations, which may be inconsistent or even conflicting. In Mitamura *et al*. ^24^, significant differences in macrophage abundance were reported between Cov and DRESS groups, as well as between Cov and NHS groups, for the Mitamura dataset. However, when using Phenograph for clustering, these differences were only observed at *k* = 30 and *k* = 100 while disappeared under other parameter settings. Additionally, CD8+ T cells were not annotated at *k* = 50 and *k* = 100. When using FlowSOM for clustering, none of the parameters that we tested could replicate the reported differences in macrophage abundance. Interestingly, at *k* = 11, CD8+ T cells showed significant abundance differences between Cov and NHS group, contradicting the original findings (Figure S2).

Our results highlight the sensitivity of unsupervised clustering methods to hyperparameter choices, and demonstrate that manual tuning of the parameter can significantly affect clustering results, leading to inconsistent results in downstream cell type identification and abundance quantification. More accurate and robust methods that perform uniformly well across different experimental conditions are desired. Consequently, unsupervised clustering methods are not included in our subsequent analyses.

### Clever enables accurate cell abundance quantification and avoids misclassifying unlabeled cell types

We evaluated the accuracy of cell classification and abundance quantification of classification based methods. Experiments were conducted on the labeled portion of the IMC dataset Hoch. We used 60% of the labeled cells in the original dataset as labeled data, and treated the remaining cells as unlabeled data. In addition, we left all the tumor cells as unlabeled cell in our test to emulate the case where certain cell types were not defined in the gating strategy in selecting the positive cell classes. Cell labels generated by manual gating were used as the ground truth for quantitative assessment. As shown in Figure 2A, compared with the three methods capable of predicting Unknown cell types, Clever demonstrated particularly high accuracy in classifying labeled positive cell types, while correctly classified the unlabeled tumor cells as “Unknown”, as illustrated in UMAP results. By contrast, CyAnno tended to overclassify a diverse range of cell types as the Unknown type. Astir predominantly categorized a significant proportion of unlabeled tumor cells as FOXP3+ T cells. The ACDC results demonstrate high accuracy in classifying unlebeled tumor cells as “Unknown”, but exhibit misclassification among positive labeled cell types, such as FOXP3+ T cells being grouped with CD4+ T cells. The other four learning methods including DGCyTOF, RandomForest, SVM and XGBoost struggled with the cells of unlabeled cell types and misclassified them as other positive labeled cell types, leading led to inflated proportions of the positive cell types to varying degrees (Figures 2B, C and S3).

**Figure 2:**
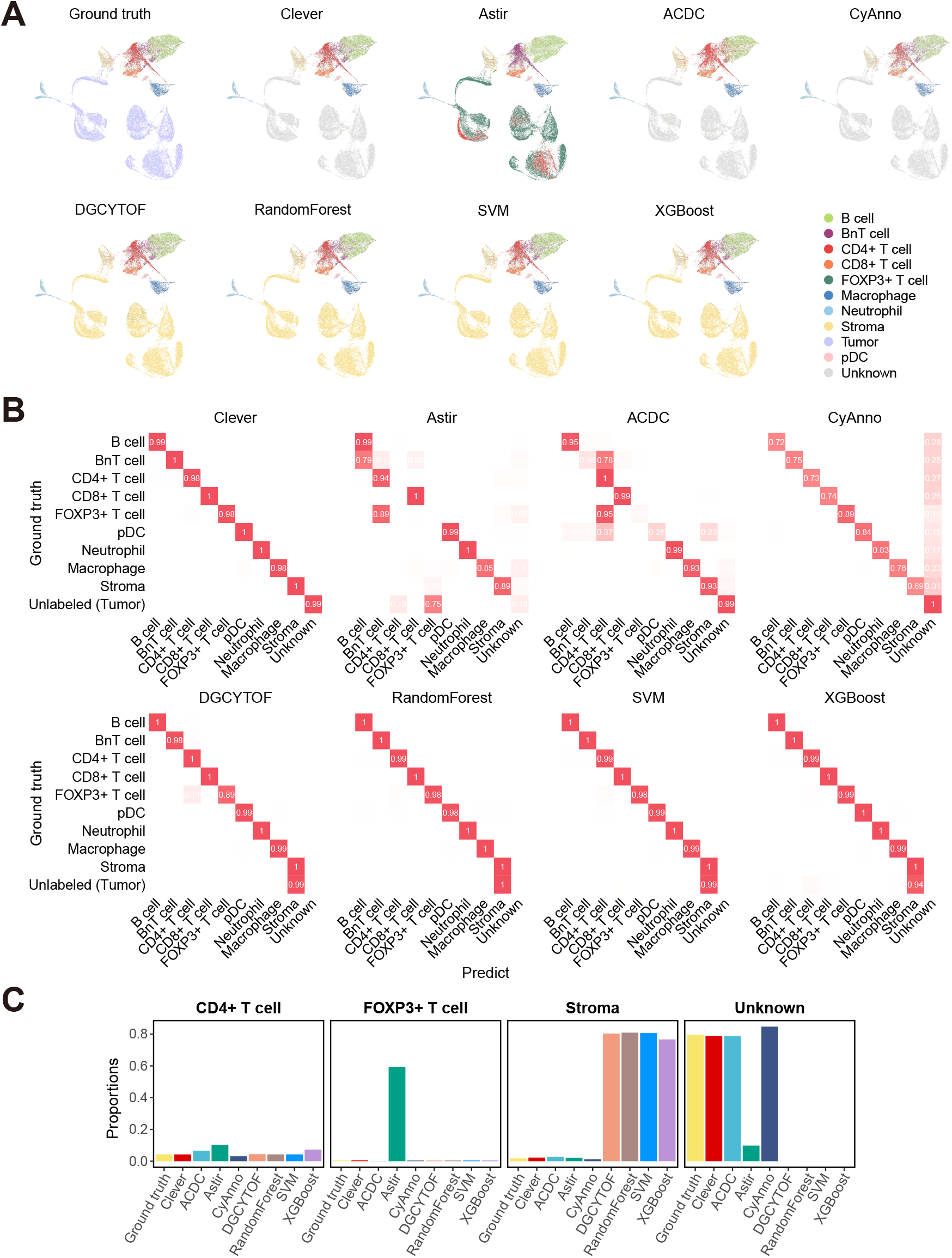
Classification and quantification performance comparison between Clever and other methods when there are unlabeled cell types in the dataset. (A) Clever successfully labels the masked tumor cells (light purple in ground truth visualization) as Unknown cell types (grey). The UMAP visualization is based on the count assay. Tumor cells, originally absent from the training set, are learned as isolated Unknown cell types. (B) Confusion matrices of Clever and other methods when tumor cells are categorized as an unseen cell type. (C) Bar charts comparing the actual and predicted abundances of specific cell types.

Next, we investigated how varying the proportions of the unlabeled tumor cells affected cell type quantification. The proportion of the tumor cells was systematically adjusted in the dataset to constitute 5%, 10%, 20%, 40%, and 60% of the total cell population. The correlations between the true proportions of tumor cells in the sample and the percentages of cells assigned to the “Unknown” type predicted by each method were shown in Figure 3A. Clever achieved the highest Lin’s CCC and the lowest RMSE among the compared methods. Note that DGCyTOF, Random Forest, SVM, and XGBoost demonstrated negative CCC and high RMSE values.

**Figure 3:**
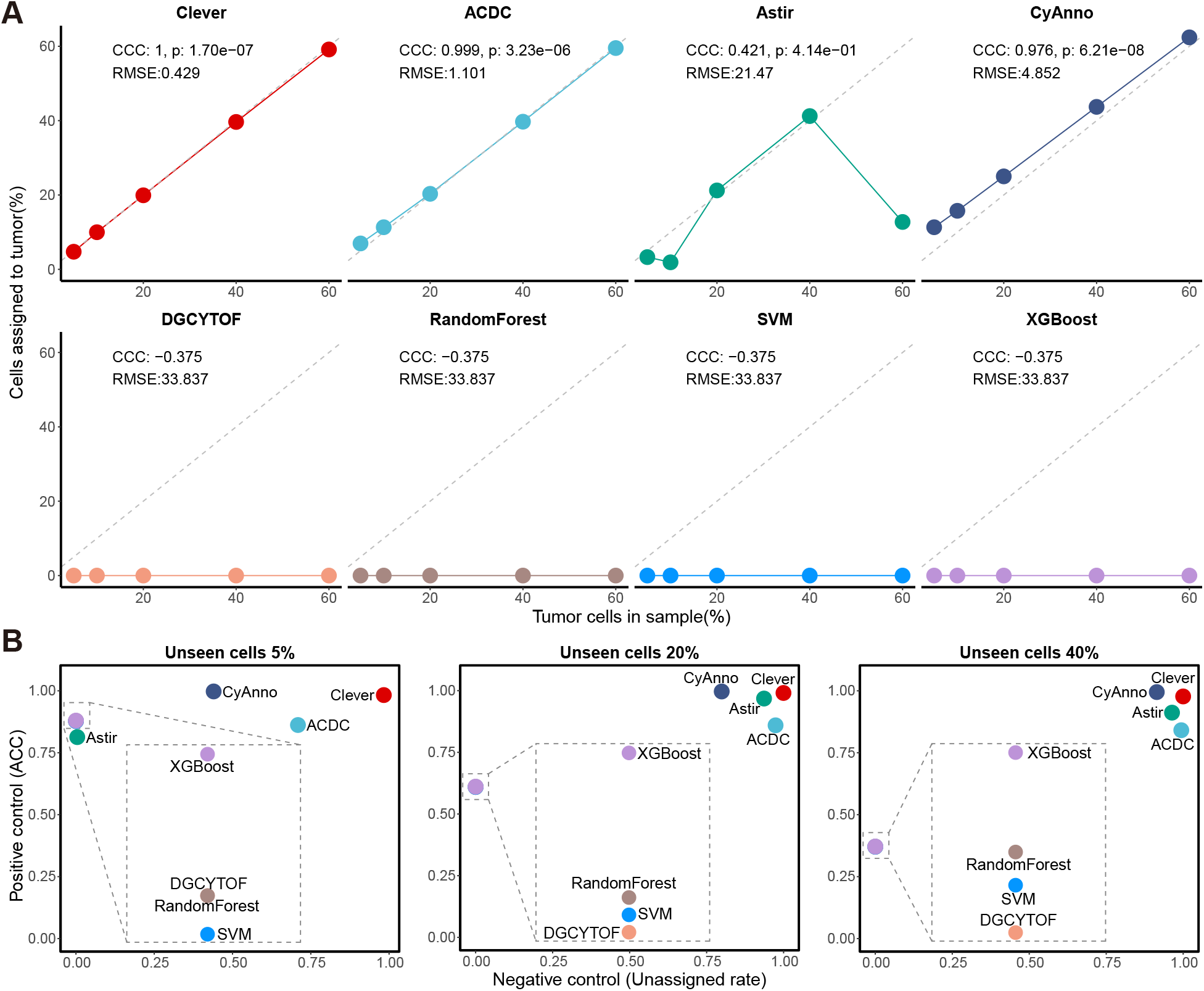
Performance evaluation with varying proportions of unseen cell types in the dataset. (A) Scatterplots of ground truth fractions (x-axis) versus predicted cell fractions (y-axis) for Clever and other methods. The plots demonstrate the predictive outcomes for the abundance of tumor cells across varying proportions. The gray dashed lines represent the *y* = *x* diagonal. Numbers within the plotting area indicate CCC and RMSE values, as well as p-values (Student’s t-test). (B) Scatterplots showing performance in positive and negative control scenarios under varying proportions of unseen cells.

In real-world application scenarios, effective cell assignment is expected to achieve reliable generalization performance on both positive controls (i.e., labeled positive cell types) and negative controls (i.e., unknown or novel cell types) to ensure accurate categorization and assignment simultaneously. To this end, we calculated the accuracy as the proportion of correctly assigned cells among labeled positive cell types, and the unassigned rate as the proportion of correctly predicted “Unknown” cells to all cells predicted as “Unknown” as performance measures on positive and negative controls, respectively. Results showed that Clever achieved the best performance in both measures (Figure 3B), while most other benchmarking methods failed to keep a balance between these two measures. The performances of DGCyTOF, random forest, SVM, and XGBoost declined significantly as the proportion of unlabeled tumor cells increased, which is as expected, since these methods are not designed to handle cases where some cell types are not included in the training set. Overall, Clever outperformed all other methods measured by a range of metrics under different proportions of unlabeled cell type in dataset (Figure S4).

Finally, we comprehensively evaluated the performance of Clever on two IMC and two CyTOF public datasets and compared its performance of cell type classification and cell abundance quantification with seven other methods. In order to simulate various positive cell labeling scenarios, we employed a leave-one-cell-type-out strategy whereby each time the labeled positive cells of a certain cell type were removed and treated as unlabeled cells. The average performances across all cell types were then compared. Clever outperformed all other methods under comparison, attaining an average accuracy of 0.958 across four datasets (Figure 4A), and demonstrating superior performance in ARI and F1-score (Figure 4B). In addition, Clever showed the lowest RMSE and the highest Lin’s CCC values across all four public datasets in cell abundance quantification evaluations (Figure 4C).

**Figure 4:**
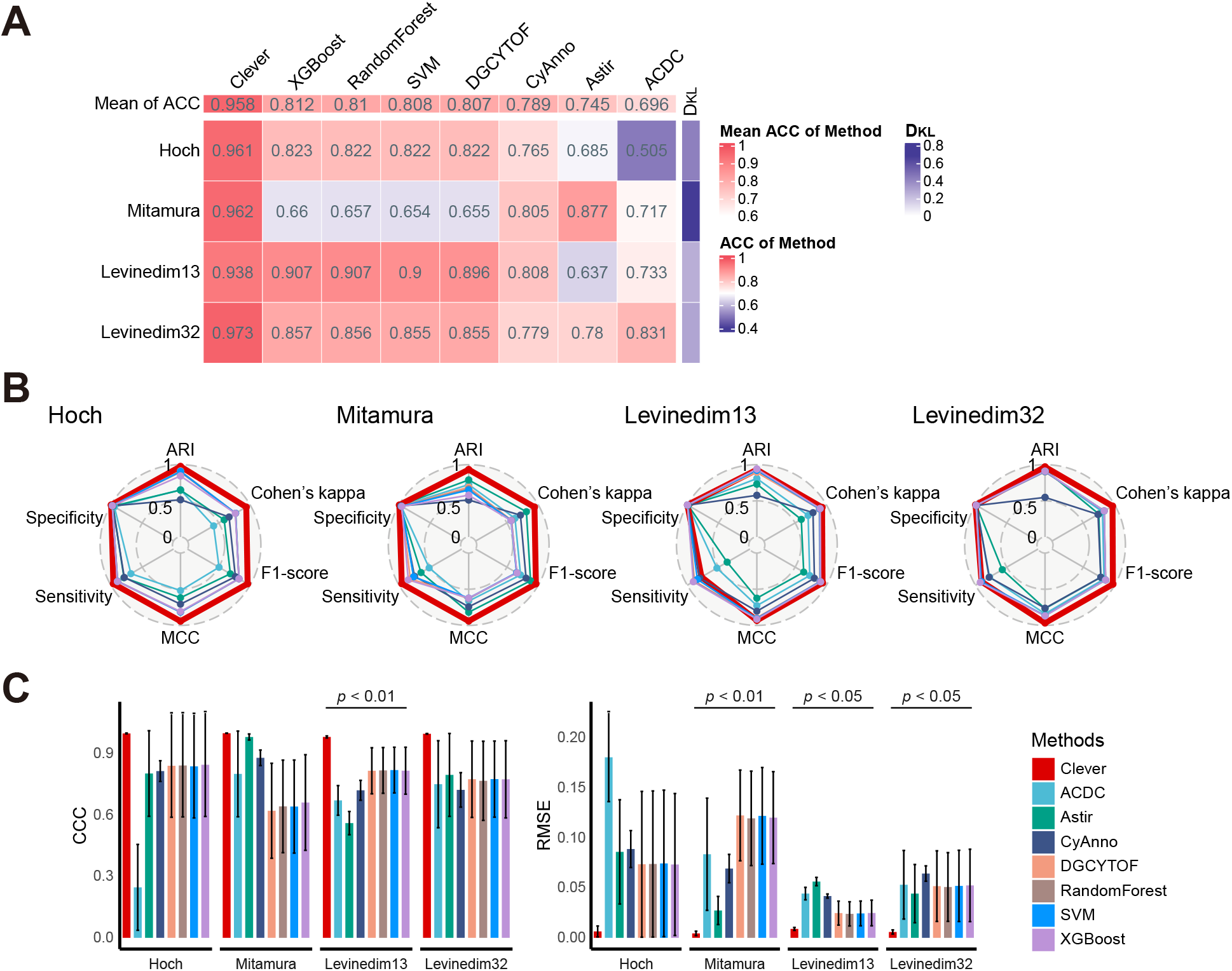
Comprehensive benchmarking between Clever and seven other methods. (A) Clever ranks first in mean accuracy compared to seven other cell type annotators across four datasets. Columns are sorted by the mean accuracy of each method on all datasets (top). The Kullback-Leibler divergence (*D*_KL_) between labeled and unlabeled data is indicated on the right. (B) Radar plots displaying the performance of Clever and other methods, evaluated using six different metrics across four datasets. (C) Boxplots showing the mean per-cell-type CCC (left) correlations and RMSE (right) for the eight methods. All data in bar plots are presented as the mean ± SD. Statistical significance is assessed using a paired t-test.

### Clever is robust to inconsistent positive cells labeling

Usually, the cell abundance results from supervised learning based cell annotation methods can be highly sensitive to the labeled positive cells used in training the cell classification models. Unfortunately, in many cases these labeled cells are identified without a consistent standard. For example, when the labeled cells are identified through manual gating, even slight adjustments to the gating boundaries can significantly alter the number of cell labels assigned. To evaluate how variation in the number of positive cell used in training impacts the performance of cell abundance quantification methods, we create new training sets containing labeled cells obtained with different marker protein expression thresholds from the Hoch dataset to emulate potential variation in manual gating process (Methods).

Figure 5B illustrates how cell abundance quantification results varies for Clever and other automated classification methods as the gating threshold changes. Clever’s quantification remains relatively stable across different gating thresholds, with minimal variation as the positive marker threshold increases. In contrast, as the gating thresholds decreases, CyAnno increasingly categorizes more cells as “Unknown”, which adversely affects the accuracy of the quantification results of known cell types. Additionally, methods such as DGCyTOF, Random Forest, and SVM display minor fluctuations in predictive performance under different gating conditions but frequently misclassify unlabeled tumor cells as stromal cells. Although these methods remain relatively stable across varying gating levels, they still face notable misclassification issues, especially with unlabeled cell types, indicating a lack of generalization in handling such cases. This observation is further supported by the MSE between the predictions of each method under different gating conditions and the original gating results. Clever achieves a low MSE under various settings of gating thresholds, indicating superior predictive stability and accuracy. In contrast, CyAnno’s MSE varies significantly with varying gating thresholds. DGCyTOF, Random Forest, SVM, and XGBoost showed higher MSE values, indicating lower predictive accuracy. These findings demonstrate that Clever maintains robustness and accuracy when the number of labeled cells varies, e.g., due to manual gating process.

**Figure 5:**
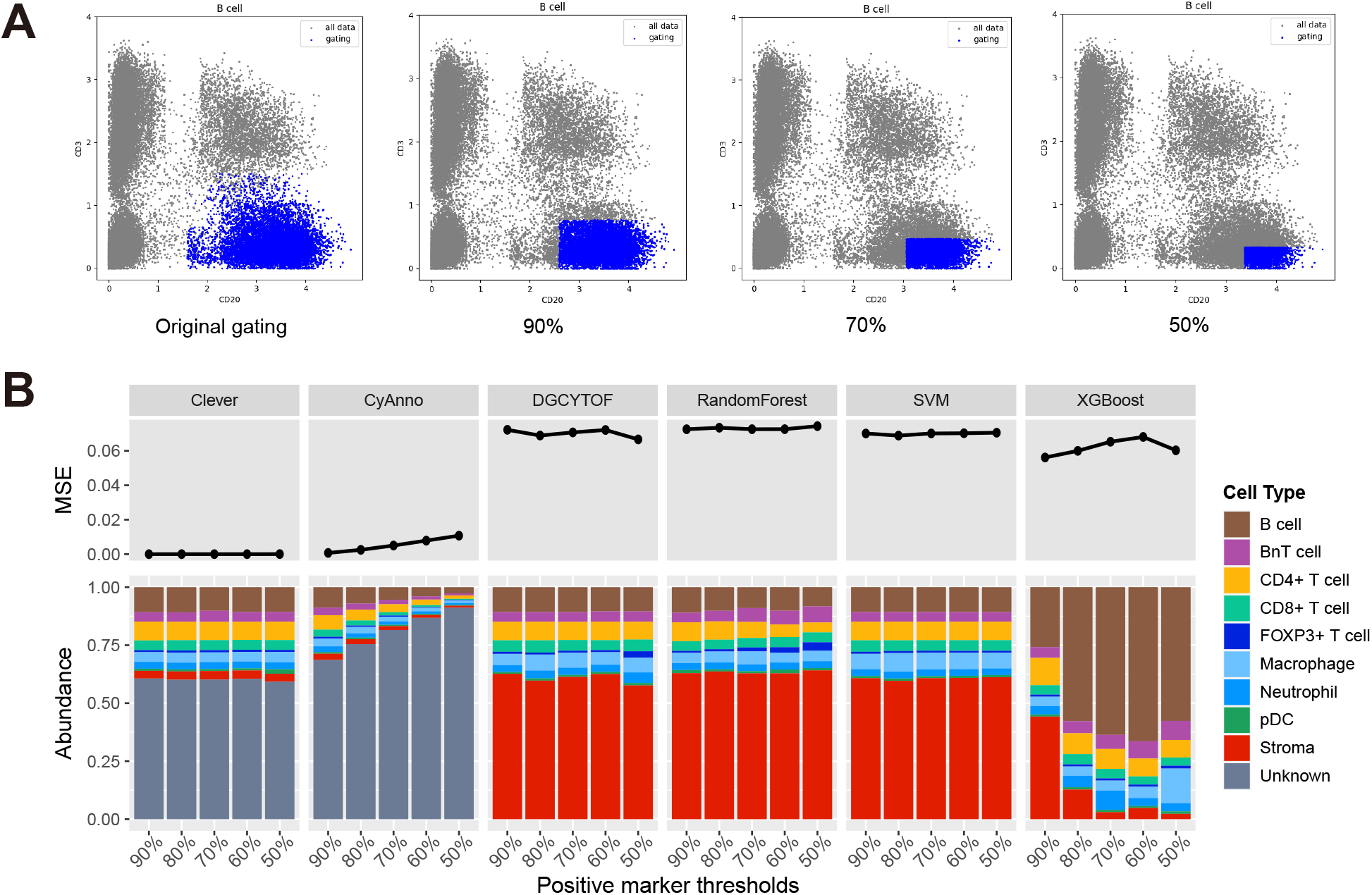
Performance of cell abundance estimation across varying gating thresholds. (A) The double-axis diagram illustrates the results of the simulated gating process. Gray dots represent all cells that have been gated, while blue dots show the re-gating results specifically for B cells under varying positive marker thresholds. (B) Comparison of mean squared error (MSE) and cell type abundance across different classification methods under different positive marker thresholds.

### Clever identifies true significant abundance differences and reduces false positives in case-control single-cell studies

We wonder if Clever and other benchmarking methods could accurately identify phenotype-relevant cell subsets with true abundance differences between groups when unlabeled cell types presented in the dataset. To this end, we generated a simulated dataset with ground truth labels (Methods) which included four distinct cell types. Cell types A and B showed no significant differences between case and control groups, while cell types C and D were designed with substantial variations between the two groups (Figure 6A). During training, only Cell type A, B and D were included in the labeled cell types and type C was not labeled.

**Figure 6:**
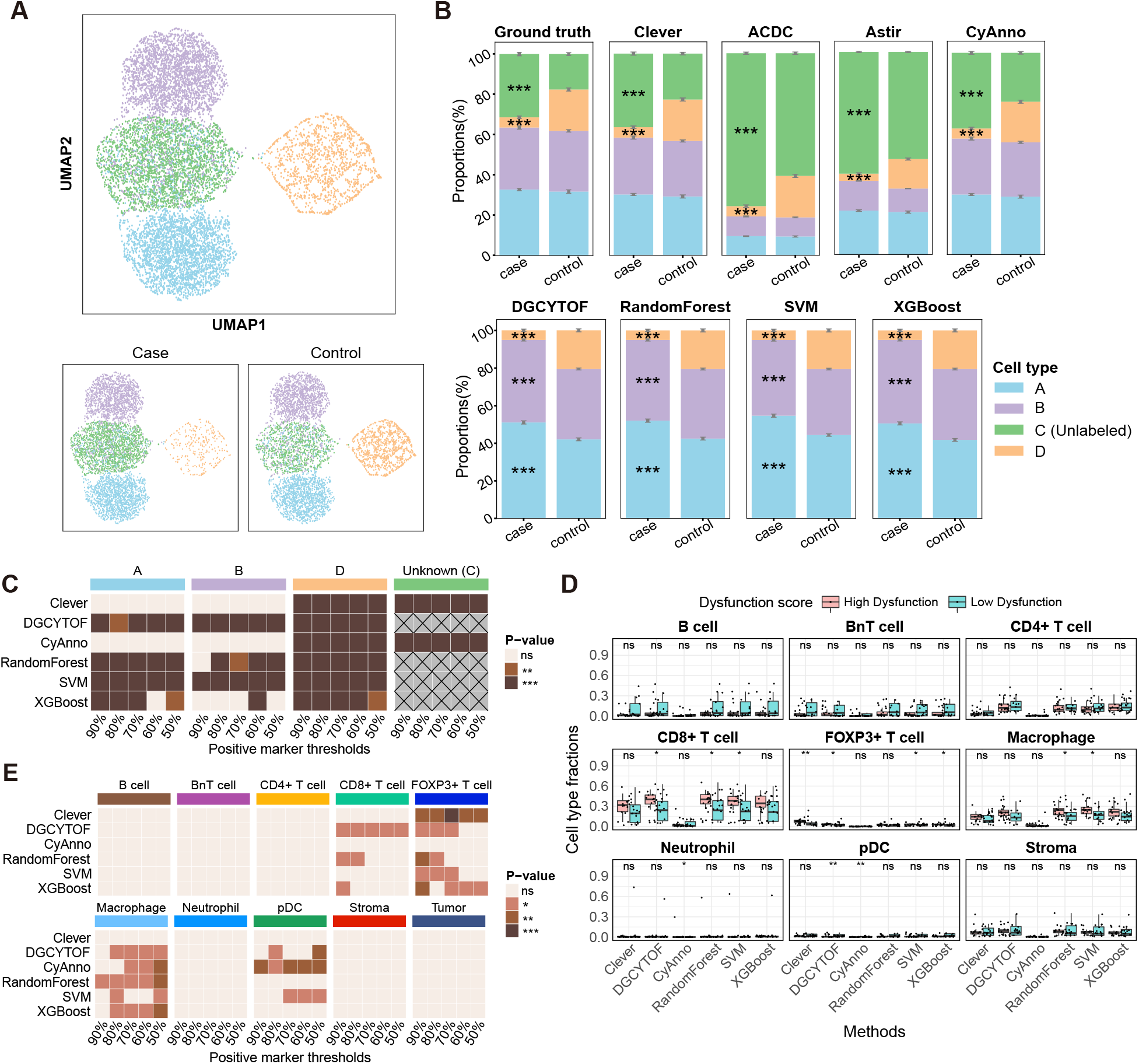
Enhanced biological accuracy and cell abundance estimation using Clever. (A) UMAP plot of the simulated dataset, displaying its two groups: case and control. This plot showcases a subset of 10,000 cells from the simulated dataset. (B) Bar plots comparing cell type fractions between case and control groups in the simulated dataset across different methods. Statistical significance is indicated by asterisks: \*\*\**p <* ± 0.001 (Wilcoxon rank sum test). Data are presented as means SD. (C) Heatmaps of statistical significance for comparisons of four cell types between case and control under various marker thresholds (Wilcoxon rank-sum test): \*\**p <* 0.01, \*\*\**p <* 0.001. (D) Boxplot comparing cell type fractions for major cell types, excluding tumor cells, between images classified as low or high dysfunctional. Significance (Wilcoxon rank-sum test) indicated by: \**p <* 0.05, \*\**p <* 0.01. (E) Heatmaps of significance for comparisons between low and high dysfunctional groups across major cell types under different marker thresholds (Wilcoxon rank-sum test): \**p <* 0.05, \*\**p <* 0.01, \*\*\**p <* 0.001.

Figure 6B shows the performance of Clever in comparison with other methods. Each bar chart represents the proportion of cell types A, B, C, and D within case and control groups, as identified by the respective algorithms against the ground truth. Clever accurately identified cell types C and D and correctly maintained consistent proportions for cell types A and B across both groups. In contrast, other methods exhibited varying performance levels. ACDC frequently over-estimated the proportion of cell type C in the control group, resulting in significant deviations from the ground truth. Astir inflated the number of “Unknown” cells and created a significant difference in cell type B between case and control groups, leading to false positive. CyAnno showed a tendency to overclassify cells as “Unknown”. The other four methods cannot identify the unlabeled cell types, leading to misclassifications of type C cells into other cell types. Consequently, these misclassifications result in erroneous detection of differences in cell types A and B between the case and control groups, where no true differences existed.

To further evaluate the robustness of Clever and other methods under varying gating thresholds, we applied different positive marker thresholds to the simulated dataset. Figure 6C and S5 illustrated how different methods perform in detecting cell type differences between the case and control groups across various gating thresholds. Both Clever and CyAnno consistently identified true differences for cell types C and D between the case and control groups across all thresholds. In contrast, while DGCYTOF and SVM showed consistent results across thresholds, they introduced false positives for cell types A and B, where no true differences existed. RandomForest and XGBoost not only misidentified differences in A and B but also showed inconsistent detection of C and D as the gating thresholds changed.

Furthermore, we extended the evaluation to real-world data. We divided samples in the Hoch dataset with high CD8+ T cell density into high-dysfunction and low-dysfunction groups based on the proportion of CD8+ T cells expressing CXCL13. Cell abundance in these samples was then quantified varying gating threshold settings using Clever and other methods. Clever’s analysis revealed a significant increase in regulatory T cells (FOXP3+ T cells) in samples classified as high-dysfunction, aligning with the characteristics of a dysfunctional immune microenvironment (Figure 6D). In contrast, other methods produced inconsistent biological conclusions as the gating boundaries were adjusted, indicating high sensitivity to gating thresholds and leading to unstable conclusions under the same conditions in these methods (Figure 6E, S6).

## Discussion

Efforts to enhance cell type quantification are crucial, particularly as single-cell technologies continue to evolve. Given the rapid pace of data generation, there is an urgent need for methods that produce more accurate and reproducible results. In this study, we present Clever, a novel computational framework for the quantification of cell types from single-cell multiplexed imaging and proteomic datasets. Clever addresses significant limitations inherent in current methods, including the lack of reproducibility in unsupervised clustering, the subjectivity and variability in marker-based methods, and the inability of supervised methods to accurately handle unseen cell types.

A key feature that distinguishes Clever is its use of an MPU learning strategy, which enables comprehensive modeling of both labeled and unlabeled data. Powered by MPU learning, Clever is able to accurately classify cells associated with predefined cell types and categorize those not associated with any known cell type into a separate category of unknown, therefore providing more reliable data for downstream biological analyses. Additionally, Clever incorporates a self-training mechanism that iteratively refines the classification of unlabeled cells, reducing reliance on manual gating and mitigating the effects of human biases and arbitrary gating thresholds. Clever’s robustness is further demonstrated in scenarios where certain cell types were deliberately obscured during training, simulating real-world scenarios where not all cell types may be identified through gating. In these cases, Clever maintains high performance, accurately identifying and quantifying cell types without being compromised by the presence of unseen cell populations. This capability is essential for studies aiming to investigate cellular heterogeneity in complex tissues and disease conditions.

Our results demonstrate that Clever consistently outperforms existing methods in both cell type identification and cell abundance quantification across various datasets, including IMC and CyTOF data. The performance of Clever in real-world applications was further validated through its ability to identify true abundance differences and reduce false positives under various conditions. For example, Clever accurately reflected the differences in cell proportions associated with different levels of T cell dysfunction in melanoma datasets, demonstrating its practical utility in clinical and research settings.

In summary, Clever represents a significant advancement in the field of single-cell data analysis, offering a robust and versatile framework for accurate cell type quantification. Its ability to handle both known and unknown cell types in a robust way makes it a valuable tool for enhancing our understanding of cellular interactions in health and disease. Future developments and refinements in Clever’s algorithms and implementation are anticipated to further expand its applicability and improve its integration with other analytical techniques, thereby advancing the field of computational biology and its contributions to biomedical research.

## Data availability

All expression datasets analyzed in this work, including accession codes and web links (if avaliable), are listed in Supplementary Table S2.

## Code availability

The source code for Clever-XMBD will be available at https://github.com/xmuyulab/Clever-XMBD.

## Acknowledgements

This work was supported by the National Natural Science Foundation of China (82388201 to J.H.).

## Authors contributions

Y.L., X.C. and Y.L. performed the computational analysis, analyzed data and generated figures. W.Y. and R.Y. supervised the research and designed the methodology. X.X. and Z.H. contributed to discussion and reviewed and edited the manuscript. All authors assisted in writing the manuscript.

## Footnotes

1 XMBD: Xiamen Big Data, a biomedical open software initiative in the National Institute for Data Science in Health and Medicine, Xiamen University, China.

