## Supplementary Information for "Accurate Cell Abundance Quantification using Multi-positive and Unlabeled Self-learning"

**Performance benchmarking against existing cell type identification methods.**

We conducted a comprehensive performance evaluation of Clever, comparing it with seven existing methods.

ACDC and Astir are marker-based cell type identification methods. ACDC is a semi-supervised approach that integrates biological prior knowledge to autonomously categorize cell types. We utilized ACDC through its official website (https://bitbucket.org/dudleylab/acdc), employing the default parameter settings. For ACDC's implementation, we constructed a marker table detailing the associations between specific markers and cell types, based on established biological knowledge. This table is instrumental in the classification process, with each marker assigned a distinct value reflecting its expression level in a given cell type: '1' denotes positive expression, '-1' denotes negative expression, and '0' denotes a lack of significant correlation.

Astir is a semi-supervised probabilistic model designed specifically for cell type discovery and classification. It integrates prior knowledge on marker proteins, sharing a conceptual resemblance with the ACDC approach. We employ Astir via its Python package, available at https://pypi.org/project/astir/. Unlike ACDC, Astir simplifies the input process by requiring only the positive genes in the marker file corresponding to each cell type. Consequently, we created a binary gene-cell type matrix for Astir's input, where '1' indicates a gene's role as a marker for a specific cell type and '0' signifies no such association.

DGCyTOF is a supervised machine learning model, specifically designed for the classification and discovery of novel cell types. It utilizes a custom neural network and a feedback loop mechanism that iteratively refines cell type labels to enhance classification accuracy. We accessed the model from its official repository at https://github.com/lijcheng12/DGCyTOF/ and set the parameters to their default configurations.

CyAnno is a semi-automatic method for cell type identification. The methodology begins by utilizing expression profiles from labeled cell groups, employing techniques such as Principal Component Analysis (PCA), kernel density estimation, and Nearest Neighborhood Approximation (NNA) to establish a landmark (LM) cell set for each group. Next, NNA is utilized to create a cell-type-specific training set, which is then employed for training one or more models dedicated to each cell type. Instructions for using CyAnno were sourced from the official website (https://github.com/abbioinfo/CyAnno), and its default parameter configurations were followed. During training, we excluded cell expression and ground truth data related to unseen cell types. These aspects were incorporated into the test set, enabling a comprehensive evaluation of the model's capabilities.

Our benchmark methods also include advanced ensemble learning techniques, including XGBoost, Random Forest, and Support Vector Machine (SVM). XGBoost, rooted in gradient boosting, was employed via the 'XGBClassifier()' from its official Python library. Random Forest and SVM were utilized through their respective Scikit-Learn Python classes, 'RandomForestClassifier' and 'SVC'.

**Confidence-based Multi-class Positive and Unlabeled (Conf-MPU) risk**

We first calculate the mean of the individual cell loss values, which we refer to as risk hereafter:

$$\mathrm{R}(f) = \sum_{i=1}^{m} \pi_i \mathrm{R}_{\mathrm{P}_i}^{+}(f) + \left(1 - \sum_{i=1}^{m} \pi_i\right) \mathrm{R}_{\mathrm{N}}^{-}(f), \tag{1}$$

where $\mathrm{R}_{\mathrm{P}_i}^{+}(f) = \mathbb{E}_{\boldsymbol{x} \sim p(\boldsymbol{x}|y=i)}[\ell(f(\boldsymbol{x}), i)]$ and $\mathrm{R}_{\mathrm{N}}^{-}(f) = \mathbb{E}_{\boldsymbol{x} \sim p(\boldsymbol{x}|y=m+1)}[\ell(f(\boldsymbol{x}), m+1)]$ represent the classification risks for the positive and unknown classes, respectively, and $\pi_i = p(y = i)$ denotes the prior probability of the $i$-th positive class. To estimate $\mathrm{R}_{\mathrm{N}}^{-}$, note that

$$\begin{aligned} & p(y = m+1)p(\boldsymbol{x} \mid y = m+1) \\ = \;& p(\boldsymbol{x}, y = m+1) \\ = \;& p(\boldsymbol{x}) - \textstyle\sum_{i=1}^{m} p(y = i)p(\boldsymbol{x} \mid y = i). \end{aligned} \tag{2}$$

Therefore we have:

$$\mathrm{R}(f) = \sum_{i=1}^{m} \pi_i \mathrm{R}_{\mathrm{P}_i}^{+}(f) + \mathrm{R}_{\mathrm{U}}^{-}(f) - \sum_{i=1}^{m} \pi_i \mathrm{R}_{\mathrm{P}_i}^{-}(f), \tag{3}$$

where $\mathrm{R}^-_{\mathrm{P}_i}(f) = \mathbb{E}_{\boldsymbol{x}\sim p_{\mathrm{P}_i}(\boldsymbol{x})}[\ell(f(\boldsymbol{x}), m+1)]$. Here, the loss function $\ell()$ is implemented using the cross-entropy function in this study.

The above modeling process relies on the assumption that the distribution of the unlabeled data aligns with the overall data distribution, *i.e.*, $p_{\mathrm{U}}(\boldsymbol{x}) = p(\boldsymbol{x})$. However, this assumption may be violated in real data, leading to potential bias in the risk estimation $\mathrm{R}^-_{\mathrm{U}}(f)$. To mitigate this bias, we adopt the approach proposed by Zhou et al. [1] to derive a Conf-MPU risk function as follows. Briefly, we first define $\lambda(\boldsymbol{x}) = p(y < m+1 \mid \boldsymbol{x})$ as the confidence score that a cell belongs to any positive class, assuming $\lambda(\boldsymbol{x}) > 0$. With this assumption, we further decompose $\mathrm{R}^-_{\mathrm{U}}(f)$ in Eq. (3) by incorporating a threshold parameter $0 < \delta \leq 1$ as follows:

$$\mathrm{R}^-_{\mathrm{U}}(f) = \sum_{i=1}^{m} \pi_i \mathrm{R}^-_{\tilde{\mathrm{P}}_i}(f) + \mathrm{R}^-_{\tilde{\mathrm{U}}}(f), \tag{4}$$

$$\mathrm{R}^-_{\tilde{\mathrm{P}}_i}(f) = \mathbb{E}_{\boldsymbol{x}\sim p_{\mathrm{P}_i}(\boldsymbol{x}|\lambda(\boldsymbol{x})>\delta)}\left[\ell(f(\boldsymbol{x}), m+1)\frac{1}{\lambda(\boldsymbol{x})}\right], \tag{5}$$

$$\mathrm{R}^-_{\tilde{\mathrm{U}}}(f) = \mathbb{E}_{\boldsymbol{x}\sim p(\boldsymbol{x}|\lambda(\boldsymbol{x})\leq\delta)}\left[\ell(f(\boldsymbol{x}), m+1)\right]. \tag{6}$$

Then, the risk function in Eq. (3) can now be rewritten as follows:

$$\mathrm{R}(f) = \sum_{i=1}^{m} \pi_i \left(\mathrm{R}^+_{\mathrm{P}_i}(f) + \mathrm{R}^-_{\tilde{\mathrm{P}}_i}(f) - \mathrm{R}^-_{\mathrm{P}_i}(f)\right) + \mathrm{R}^-_{\tilde{\mathrm{U}}}(f), \tag{7}$$

where $\mathrm{R}^+_{\mathrm{P}_i}(f)$ and $\mathrm{R}^-_{\mathrm{P}_i}(f)$ calculate the classification risk for labeled positives, and $\mathrm{R}^-_{\tilde{\mathrm{P}}_i}(f)$ and $\mathrm{R}^-_{\tilde{\mathrm{U}}}(f)$ calculate the classification risk for the high-confidence positive and unknown classes within the training data, respectively.

In Conf-MPU, given an appropriate threshold $\delta$, $\lambda(\boldsymbol{x}) > \delta$ indicates that $\boldsymbol{x}$ is a positive sample; otherwise, it is considered a unknown sample. This implies that $p_{\mathrm{P}_i}(\boldsymbol{x} \mid \lambda(\boldsymbol{x}) > \delta) \approx p_{\mathrm{P}_i}(\boldsymbol{x})$, and $p(\boldsymbol{x} \mid \lambda(\boldsymbol{x}) \leq \delta) \approx p_{\mathrm{U}}(\boldsymbol{x} \mid \lambda(\boldsymbol{x}) \leq \delta) \approx p_{\mathrm{N}}(\boldsymbol{x})$, even if $p_{\mathrm{U}}(\boldsymbol{x})$ differs from $p(\boldsymbol{x})$. Thus, $\mathrm{R}^-_{\tilde{\mathrm{P}}_i}(f)$ and $\mathrm{R}^-_{\tilde{\mathrm{U}}}(f)$ can be estimated with reduced bias using $\mathcal{X}_{\mathrm{P}_i}$ and $\mathcal{X}_{\mathrm{U}}$, respectively. This approach leads to a more accurate estimation of $\mathrm{R}^-_{\mathrm{U}}(f)$. Based on the foregoing deductions, the risk estimator can be formulated as follows:

$$
\begin{aligned}
\mathrm{R}_{\text{Conf-MPU}}(f) =& \boldsymbol{w}_p \left( \sum_{i=1}^{m} \frac{\pi_i}{n_{\mathrm{P}_i}} \sum_{j=1}^{n_{\mathrm{P}_i}} \max\{0, \ell(f(\boldsymbol{x}_j^{\mathrm{P}_i}), i) \right. \\
& \left. + \mathbb{1}_{\hat{\lambda}(\boldsymbol{x}_j^{\mathrm{P}_i}) > \delta} \ell(f(\boldsymbol{x}_j^{\mathrm{P}_i}), m+1) \frac{1}{\hat{\lambda}(\boldsymbol{x}_j^{\mathrm{P}_i})} - \ell(f(\boldsymbol{x}_j^{\mathrm{P}_i}), m+1)\} \right) \\
& + \frac{1}{n_{\mathrm{U}}} \sum_{j=1}^{n_{\mathrm{U}}} \mathbb{1}_{\hat{\lambda}(\boldsymbol{x}_j^{\mathrm{U}}) \leq \delta} \ell(f(\boldsymbol{x}_j^{\mathrm{U}}), m+1),
\end{aligned} \tag{8}
$$

where $\hat{\lambda}$ serves as an empirical confidence score estimator. In our model, a classifier with a sigmoid output layer fulfills this role, ensuring that $\hat{\lambda}(\boldsymbol{x}) > 0$. Furthermore, we determine the empirical probability $\pi_i$ based on the proportion of each positive class within individual training batches. We incorporate $\mathrm{R}^+_{\mathrm{P}_i}(f)$, $\mathrm{R}^-_{\tilde{\mathrm{P}}_i}(f)$ and $\mathrm{R}^-_{\mathrm{P}_i}(f)$ in Eq. (7) as components contributing to the positive risk. Each of these risks is assigned a weight, denoted as $\boldsymbol{w}_p$, to control the importance on positive versus unknown classes.


1. Zhou, K., Li, Y. & Li, Q. Distantly Supervised Named Entity Recognition via Confidence-Based Multi-Class Positive and Unlabeled Learning. In

Proceedings of the 60th Annual Meeting of the Association for Computational Linguistics, 7198–7211 (2022).

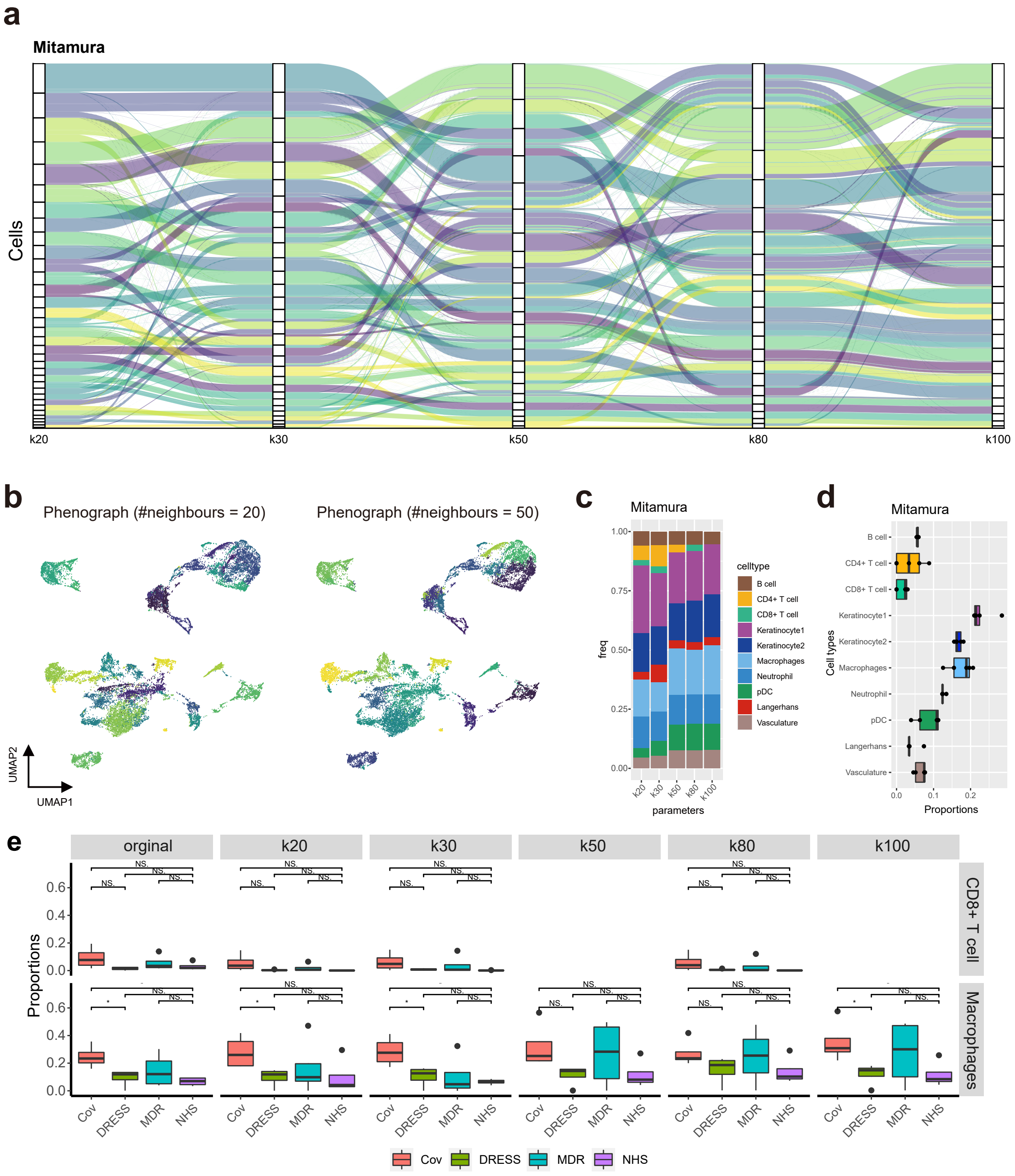


Figure S1: **Performance of Phenograph clustering results under different hyperparameters on the "Mitumura" dataset.** (a) Sankey plot comparing Phenograph clustering results with different hyperparameters for "Mitamura" dataset. (b) UMAP visualizations of "Mitamura" dataset, colored by unsupervised clustering results. The hyperparameters corresponding to the number of neighbors were set to 20 and 50. (c) Stacked bar charts showing cell percentages, colored by 10 main cell types. (d) Boxplots displaying the distribution of cell percentages with different hyperparameters for each cell type. (e) Boxplots illustrating the proportions of Macrophages and CD8+ T cells between samples with different symptoms.

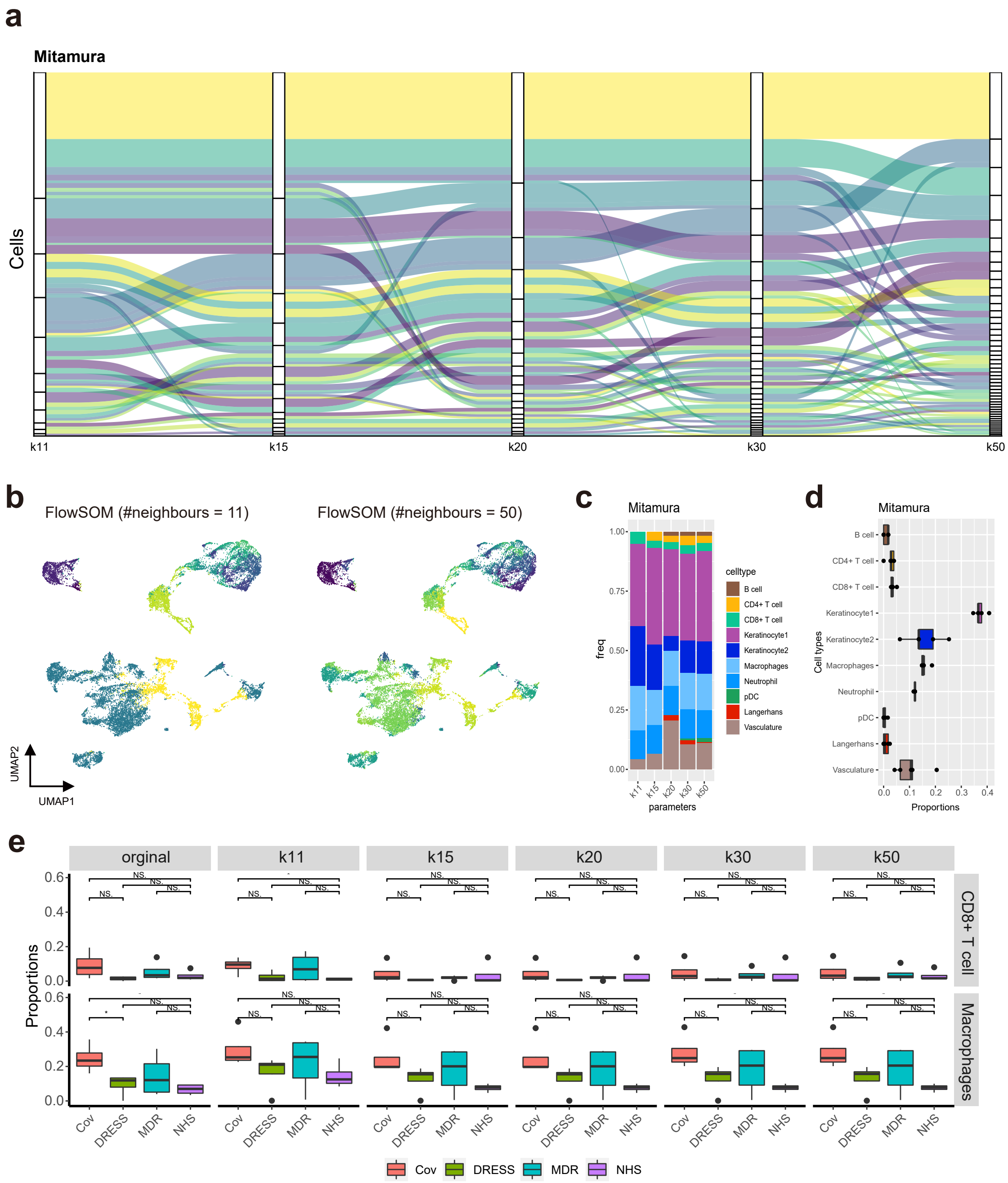


Figure S2: **Performance of FlowSOM clustering results under different hyperparameters on the "Mitumura" dataset.** (a) Sankey plot comparing FlowSOM clustering results with different hyperparameters for "Mitamura" dataset. (b) UMAP visualizations of "Mitamura" dataset, colored by unsupervised clustering results. The hyperparameters corresponding to the number of neighbors were set to 11 and 50. (c) Stacked bar charts showing cell percentages, colored by 10 main cell types. (d) Boxplots showing the distribution of cell percentages with different hyperparameters for each cell type. (e) Boxplots showing the proportions of Macrophages and CD8+ T cells between samples with different symptoms. The significance of the two-sided T-test is represented by stars: $*p < 0.05$, $**p < 0.01$, $***p < 0.001$. The abbreviation "ns" indicates a $p$-value greater than 0.05

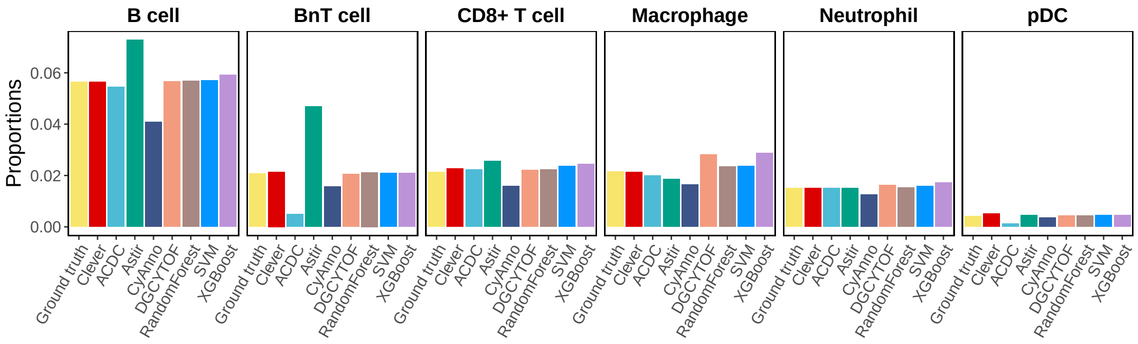


Figure S3: Proportions of each cell type in the ground truth and those predicted by different methods. The results are shown for the “Hoch” query dataset with Tumor as the unseen cell type.

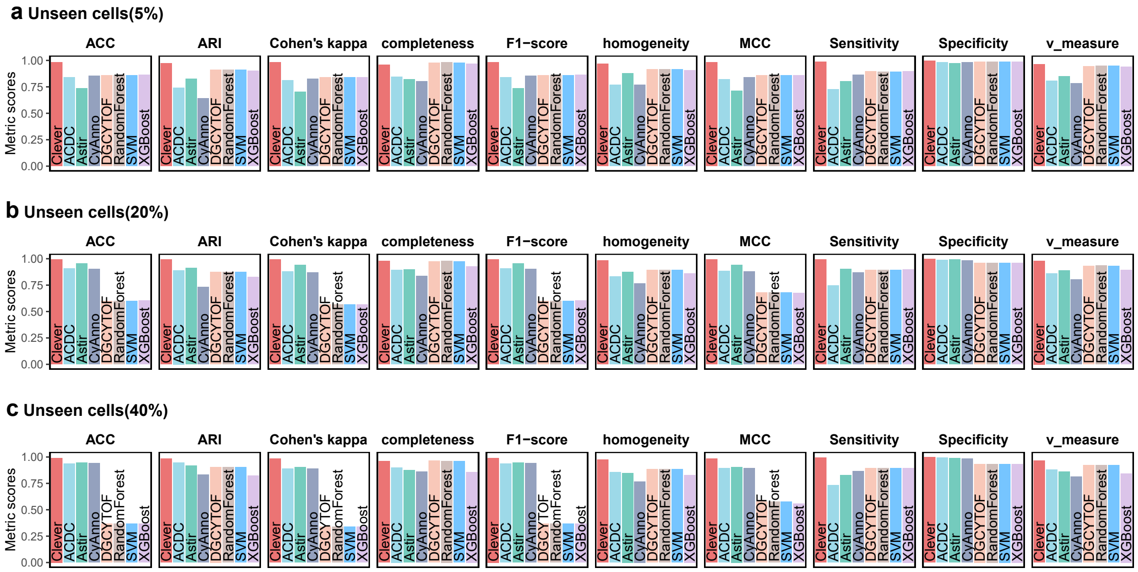


Figure S4: Performance evaluation with varying proportions of unseen cell types in the Hoch dataset: (a) 5%, (b) 20%, (c) 40%. Bar charts illustrate the performance of each method across different metrics.

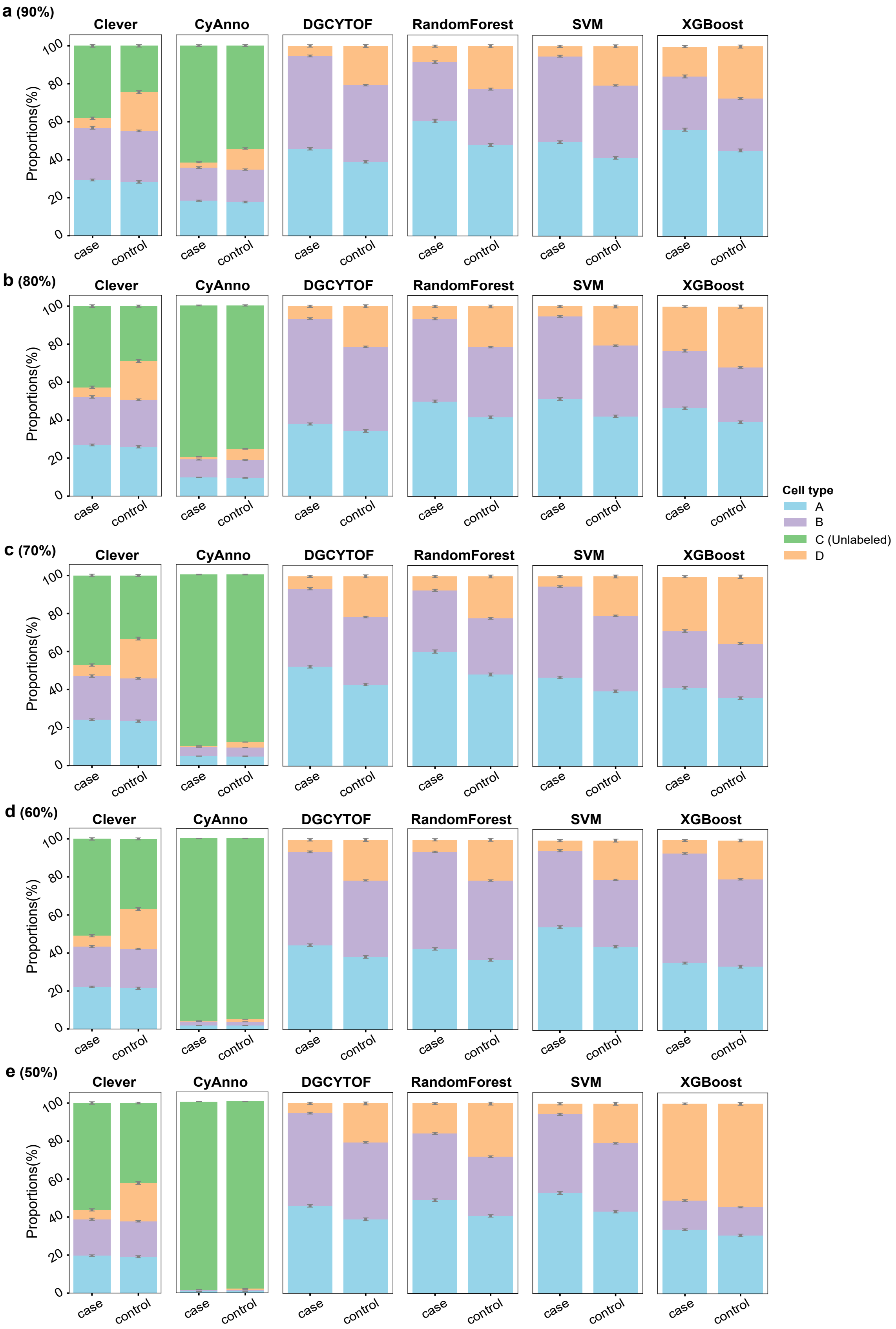


Figure S5: Bar plots comparing cell type fractions between case and control groups in the simulated dataset using different methods under various positive marker thresholds: (a) 90%, (b) 80%, (c) 70%, (d) 60%, (e) 50%.

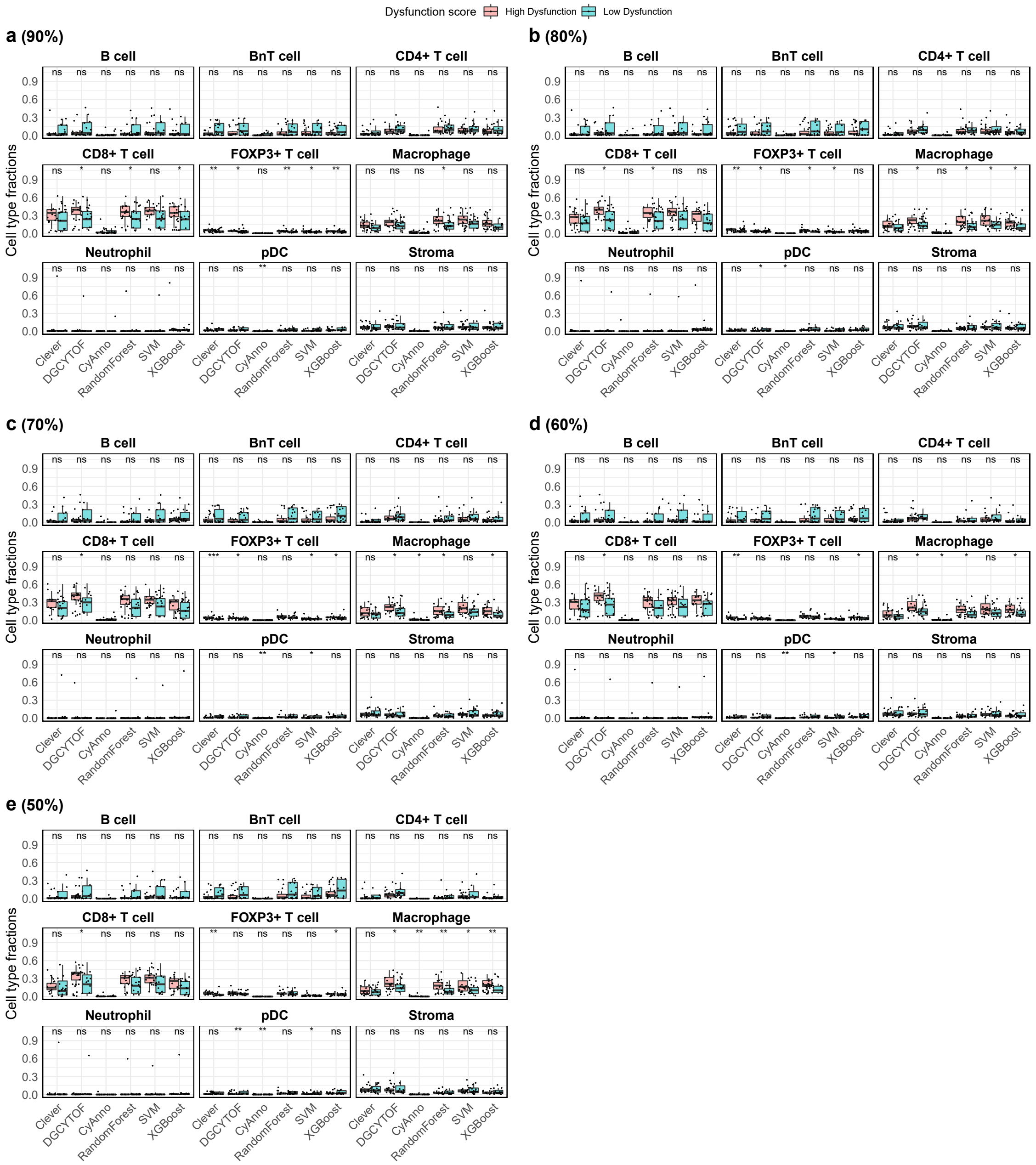


Figure S6: Boxplot comparing the fractions of major cell types, excluding tumor cells, between images classified as low or high dysfunction under different positive marker thresholds: (a) 90%, (b) 80%, (c) 70%, (d) 60%, (e) 50%. Statistical significance between groups is indicated by asterisks: $*p < 0.05$, $**p < 0.01$ (Wilcoxon rank sum test).

### Supplementary Tables

**Table S1:** Overview of related methods for cell type identification.

| Methods | Functions | Source | Link | Reference |
|---|---|---|---|---|
| FlowSOM | Unsupervised clustering algorithm | R package FlowSOM; Cytofkit | https://github.com/JinmiaoChenLab/cytofkit | Van Gassen et al. (2015) |
| Phenograph | Unsupervised clustering algorithm | R package Rphenograph; Cytofkit | https://github.com/JinmiaoChenLab/cytofkit | Levine et al. (2015) |
| ACDC | Marker-based automated cell type identification | Python script | https://bitbucket.org/dudleylab/acdc/src/master | Lee et al. (2017) |
| Astir | Marker-based automated cell type identification | Python script | https://pypi.org/project/astir/ | Geuenich et al. (2021) |
| CyAanno | Semi-automated ML-based computational framework | Python script | https://github.com/abbioinfo/CyAnno | Kaushik et al. (2021) |
| DGCyTOF | Supervised classification | Python script | https://github.com/ lijcheng12/DGCyTOF/ | Cheng et al. (2022) |
| RandomForest | Supervised classification | Python script | https://github.com/scikit-learn/scikit-learn | Breiman (2001) |
| SVM | Supervised classification | Python script | https://github.com/scikit-learn/scikit-learn | Cortes and Vapnik (1995) |
| XGBoost | Supervised classification | Python script | https://github.com/scikit-learn/scikit-learn | Chen and Guestrin, (2016) |

**Table S2:** Inventory of expression datasets analyzed in this work.

| Database | No. of markers | No. of manually gated populations | No. of manually gated cells (labeled data) | Source | Reference |
|---|---|---|---|---|---|
| Hoch | 46 | 11 | 87,696 | https://doi.org/10.5281/zenodo.5994136 | Hoch et al. (2022) |
| Mitamura | 39 | 13 | 22,854 | https://doi.org/10.5281/zenodo.5036924 | Mitamura et al. (2021) |
| Levinedim 13 | 13 | 24 | 81,747 | https://github.com/lmweber/benchmark-data-Levine-13-dim | Bendall et al. (2011) |
| Levinedim 32 | 32 | 14 | 104,184 | https://github.com/lmweber/benchmark-data-Levine-32-dim | Levine et al. (2015) |
